# Visual search for upright bigrams predicts reading fluency in children

**DOI:** 10.1101/2021.04.14.439823

**Authors:** Aakash Agrawal, Sonali Nag, K.V.S. Hari, S. P. Arun

**Author notes:** Correspondence to: S. P. Arun.

## Abstract

Fluent reading is an important milestone in education, but we lack a clear understanding of why children vary so widely in attaining this milestone. Language-related factors such as rapid automatized naming (RAN) and phonological awareness have been identified as important factors that influence reading fluency. Of theoretical interest is also, however, whether aspects of visual processing influence reading fluency. To investigate this issue, we tested primary school children (n = 68) on four tasks: two reading fluency tasks (word reading and passage reading), a RAN task to measure naming speed, and a visual search task using letters and bigrams to measure visual processing. As expected, the RAN score was strongly correlated with reading fluency. In addition, visual processing of bigrams was correlated with reading fluency. Importantly, this association was specific to upright but not inverted bigrams, and to bigrams with normal but not large letter spacing. Thus, reading fluency in children is accompanied by specialized changes in upright bigram processing. We propose that bigram processing during visual search could complement existing measures of language processing to understand individual differences in reading fluency.

## INTRODUCTION

Learning to read fluently is an important milestone during development, but there is considerable individual variation in attainment. For alphabetic languages, this variation has been explained using two simpler cognitive tasks: phoneme awareness (PA, which measures the ability to manipulate phonemes in a word), and rapid automatized naming (RAN, which measures the speed of naming visually presented letters or objects) (Melby-Lervåg et al., 2012; Norton and Wolf, 2012). These abilities not only explain concurrent individual variation in reading fluency (Melby-Lervåg et al., 2012; Norton and Wolf, 2012), but also its longitudinal development (Parrila et al., 2004; Lervåg and Hulme, 2009; Landerl et al., 2018; Vander Stappen and Reybroeck, 2018).

The RAN measure has been hypothesized to capture efficiency in cross-modal print processing (Nag and Snowling, 2012). Other explanations for the robust RAN-reading association range from domain-general speed of processing (Kail et al., 1999), especially serial processing (Sideridis et al., 2016), to domain-specific speed to retrieve phonological codes, discriminate component visual features (Stainthorp et al., 2010) and recognize whole visual items (Lervåg and Hulme, 2009). Thus, RAN captures component processes that are both perceptual-lexical as well as attentional and memory-based (Sideridis et al., 2016).

Given that reading begins with vision, it stands to reason that fluent reading is associated with changes in visual processing as well as in phonological or naming abilities. However, most previous work has focused on attentional deficits, particularly with respect to reading difficulties. Dyslexia is associated with a range of processing deficits in visuospatial attention (Goswami, 2015), crowding (Bouma and Legein, 1977; Martelli et al., 2009; Zorzi et al., 2012), attention span (Bosse et al., 2007), change detection (Rima et al., 2020) and visual search (Casco and Prunetti, 1996; Vidyasagar and Pammer, 1999). Whether these deficits explain normal variation in reading skills is, however, not clear. At a more basic level, it is not clear whether visual representations of letters or strings themselves change with reading experience, and whether these changes predict reading fluency.

It is widely believed that learning to read leads to the formation of specialized detectors for letter combinations (Grainger and Whitney, 2004; Dehaene et al., 2005). Evidence in favour of this account comes from the greater activation of the word form regions to strings containing frequent bigrams. However, recent evidence has challenged this possibility by showing that discrimination between longer strings can be explained using single letters (Agrawal et al., 2019, 2020), and that fluent readers experience weaker interactions between letters in a bigram (Agrawal et al., 2019). However this association between bigram processing and reading fluency may be explained by other factors not tested in previous studies.

### Overview of this study

Here, we investigated the relation between reading fluency and visual processing by testing two specific hypotheses. First, we asked whether learning to read results in the formation of specialized bigram detectors. Since reading involves extensive experience with upright letters, we hypothesized that learning to read would result in the formation of specialized detectors for upright bigrams but not inverted bigrams. This comparison avoids any confounds due to letter shape. To detect the presence of bigram detectors, we formulated a quantitative “letter sum” model to predict visual search on bigrams using the constituent single letters. Since bigram detectors, by definition, are activated by the entire bigram but not by the constituent letters, their presence should lead to poor performance of the letter sum model. We therefore predicted that the presence of upright bigram detectors should lead to poor performance of the letter-sum model for upright but not inverted bigrams. Comparing upright and inverted bigrams also avoids any indirect confounds due to covarying cognitive factors. For instance, a correlation between visual search performance and reading fluency could simply be due to the requirement for visuospatial attention in both tasks (Franceschini et al., 2012).

Second, we hypothesized that reading fluency variations across children would be predicted by upright bigram processing during visual search, over and above the variation predicted by RAN tasks. This is a non-trivial outcome because it implies that changes in visual processing are independent of the perceptual-lexical processes captured by RAN, and that both influence reading fluency. Alternatively, it could be that bigram processing does not predict reading fluency variations any more than RAN measures, suggesting that changes in visual processing do not directly influence reading fluency.

To assess these possibilities, we tested children in grades 3-5 (7-11 years old) across two time points (separated by ~10 months). Each child was tested on two standardized measures of reading fluency (word and paragraph reading). To reduce testing time with children, we selected a RAN task over a phoneme awareness (PA) task because the former is a better predictor of reading in some alphabetic orthographies (Landerl et al., 2018; Vander Stappen and Reybroeck, 2018), and PA is prone to floor effects in India (the location of the present study) where literacy instruction privileges either the look-and-see method or the syllable units in a word (Nag, 2017). To measure visual processing, each child was tested on a visual search task involving both single letters as well as upright and inverted bigrams. We chose visual search because it is a natural, intuitive task for children (they have to simply search for an odd-one-out), yet it has an objective measure (correctly identifying the target). At the same time, measures of search time in visual search can yield many insights into the underlying representations of visual features, including printed letters (Arun, 2012; Mohan and Arun, 2012; Pramod and Arun, 2016; Agrawal et al., 2019).

## RESULTS

Our goal was to investigate whether reading fluency can be linked to visual processing of bigrams. We selected children in grades 3-5 (aged 7-11 years) from a school where English is the medium of instruction, and tested them on English letters and words. Participants performed three tasks related to their reading skills: a word reading task (Figure 1A), a passage reading task (Figure 1B), and a rapid automatized naming (RAN) task (see Methods). As expected, the passage and word reading fluency scores were highly correlated with each other (Figure 1C). These children were further tested on a visual search task to characterize their visual processing (Figure 1D). In the visual search task, children were asked to identify an oddball target among multiple identical items.

**Figure 1.**
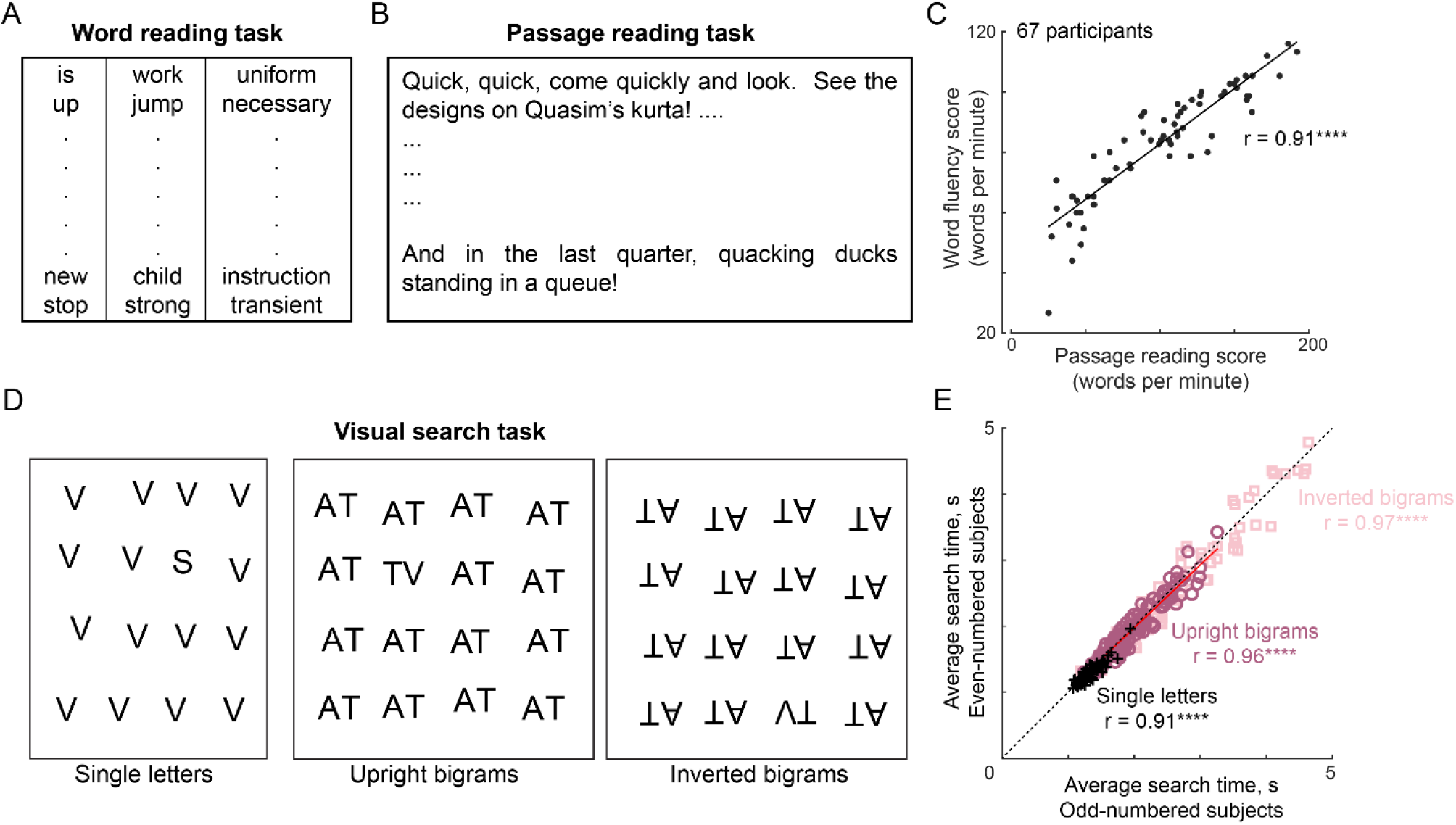
Reading fluency and visual processing tasks (Experiment 1) (A) Example words from the standardized sight word efficiency task (TOWRE). (B) The passage shown to the children to measure their reading fluency (see Methods). (C) Correlation between the fluency scores obtained from word reading task (A), and passage reading task (B). Each point represent one subject (n = 67) and asterisks indicate that the correlation is significant (**** is p < 0.00005). (D) Example single letter and bigram search array from the visual search task. (E) Split-half consistency of the visual search data for letters (+), upright bigrams (o), and inverted bigrams (□), as estimated by the correlation between search time averaged across the odd-numbered subjects and even-numbered subjects.

### Experiment 1: Single letter and bigram searches

In Experiment 1, we tested 68 children from grades 3-5 (7-11 years old) on reading tasks as described above and a visual search task. In the visual search task, both the oddball and the distractors were either single letters, or upright bigrams or inverted bigrams, and were analysed separately.

### Visual search for single letters

We first analysed the performance of the participants on single letter searches. An example search involving single letters is shown in Figure 1D. Participants were highly accurate in their performance (average accuracy across 78 single letter searches, mean ± std: 98% ± 2.4% across 68 children). They also made highly consistent responses, as evidenced by a strong and significant correlation between the average search times of odd and even-numbered participants (Figure 1E). We did not observe any significant correlation between mean single letter search time and passage reading score (r = −0.2, p = 0.1).

### Visual search for upright vs inverted bigrams

Next we sought to evaluate whether bigram processing is different for upright compared to inverted bigrams. Specifically, we reasoned that, if learning to read upright letters leads to the formation of upright bigram detectors, any model based on single letters would perform poorly on predicting upright but not inverted bigrams.

Participants performed oddball visual search in which both target and distractors were either upright or inverted bigrams (Figure 1D). As before, irrespective of fluency level, they were highly accurate in all conditions (average accuracy across 115 bigram searches, mean ± sem: 95.8% ± 0.5% for upright bigrams, 95% ± 0.7% for inverted bigrams) and also highly consistent in their responses (Figure 1E). Interestingly, participants took longer to perform inverted searches (average response times, mean ± sem across participants: 1.96 ± 0.03 s for upright, 2.43 ± 0.05 s for inverted; p < 0.00005, paired t-test across 115 searches). Thus, familiarity with the upright orientation improved discrimination. However, familiarity did not qualitatively alter visual search performance, as evidenced by a strong and significant correlation between search dissimilarities in the upright and inverted conditions (r = 0.92 across 115 bigram searches, p < 0.00005).

As with single letter analysis, we correlated the mean search time with passage reading score. Interestingly, the association between reading fluency and visual search times was specific to upright but not inverted bigrams (correlation between passage reading score and mean bigram search time: r = −0.32, p < 0.05 for upright bigrams, and r = −0.15, p = 0.24 for inverted bigrams).

### Can bigram search be explained using single letter relations?

The above findings show that reading fluency is associated with upright bigram searches, but does not elucidate whether this is due to improved single letter representations or due to specialized bigram detectors. To investigate this issue, we devised a quantitative model to explain visual search for bigrams using the constituent single letters. In a series of previous studies, we have shown that the reciprocal of search time (1/RT) – which is a measure of dissimilarity – yields more accurate models for visual search, and that the dissimilarity between objects differing in multiple features can be explained using the constituent features.

In keeping with these findings, we devised a “letter-sum” model (Figure 2A) in which the search dissimilarity (1/RT) between a pair of bigrams, say AB & CD, is a linear sum of dissimilarities between the constituent pairs of single letters A, B, C, D i.e. (A,B), (A,C), (A,D), (B,C), (B,D), and (C,D). To account for possible differences in position, we grouped these pairs based upon the type of comparison: there were letter pairs at corresponding locations in the two bigrams (e.g. AC & BD), at opposite or across locations (e.g. AD & BC), and within a bigram (e.g. AB & CD). Thus, the search dissimilarity for bigrams AB & CD is given by:

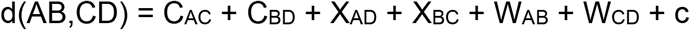

where C_AC_ & C_BD_ are relations between letters in the two bigrams at corresponding locations, X_AD_ & X_BC_ are relations between letters in the two bigrams at opposite locations, W_AB_ & W_CD_ are letter relations within each bigram and c is a constant term. The part sum model works because the same terms repeat across searches: for instance, the term C_AC_ is also present in the equation for d(AE,CF), d(AG,CH) etc. Since bigrams were constructed using six possible letters, the corresponding-location letter terms are ^6^C_2_ = 15 in number, and likewise there are 15 across-location letter terms and 15 within-bigram letter terms. These unknown part relations can then be estimated from the data using standard linear regression (see Methods).

**Figure 2.**
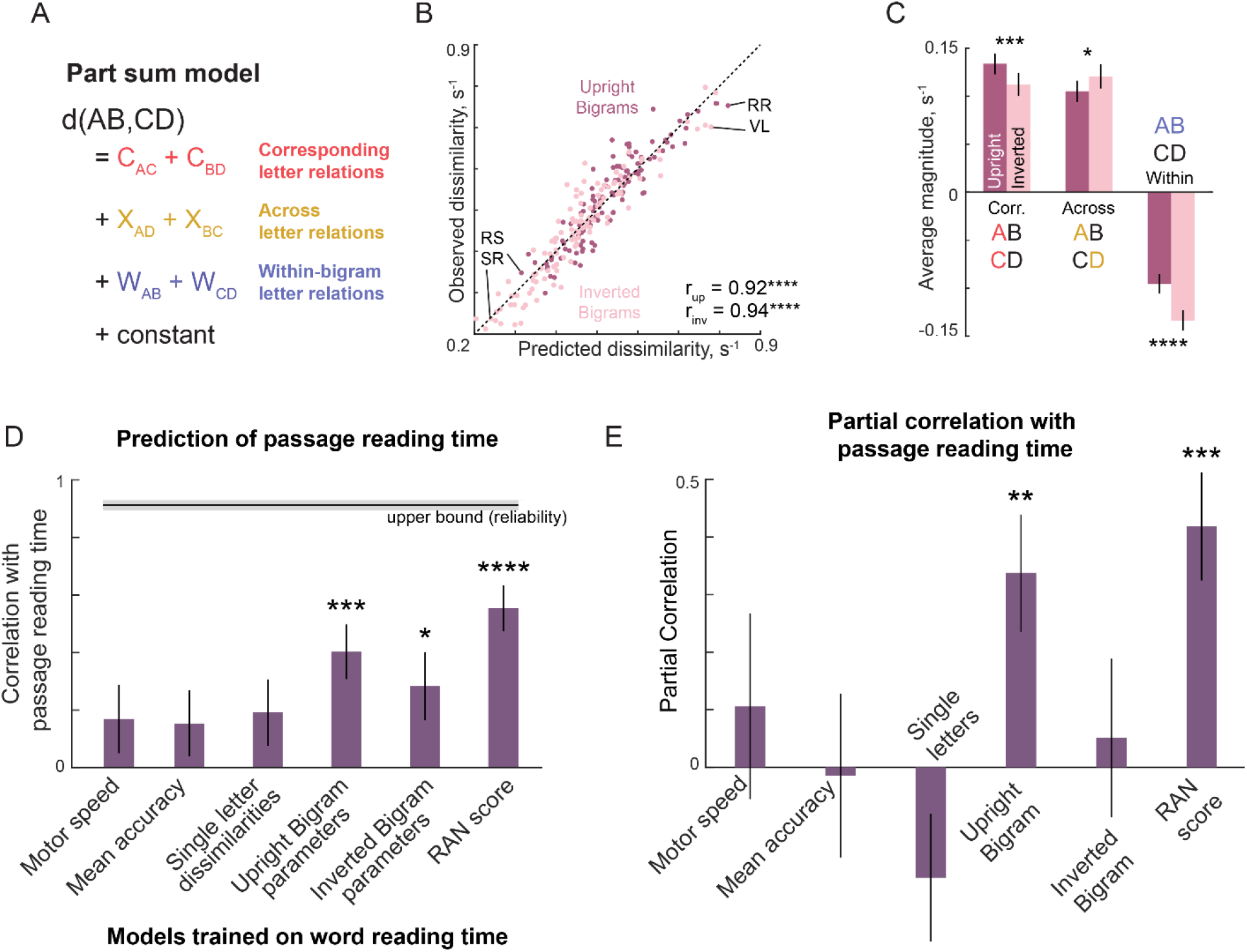
Upright bigram processing predicts reading fluency (Experiment 1) (A) Schematic of the letter-sum model, in which the net dissimilarity between two bigrams is a linear sum of single letter relations at corresponding locations across bigrams (C), opposite locations across bigrams (X) and within-bigrams (W). (B) Observed bigram dissimilarity is plotted against predicted bigram dissimilarity from the part-sum model for both upright (*dark*) and inverted (*light*) bigram searches. Each point represents one search pair (n = 115 each) and few example searches are highlighted. Asterisks indicate that the model predictions were significantly correlated with the observed dissimilarity values (p < 0.00005). (C) Average model coefficients (mean ± sem) of each type for upright and inverted bigrams. Asterisks denote statistical significance obtained on a sign-rank test comparing 15 letter dissimilarities between upright and inverted conditions (* is p < 0.05, ** is p < 0.005, etc). (D) Model correlation of each factor in predicting passage reading score. Error bars indicate ±1 s.d. using a bootstrap procedure (in which we repeatedly sampled 67 participants with replacement for a total of 1,000 times). All models were trained on word reading score, and tested on passage reading scores. Shaded error bars represent the noise ceiling i.e. correlation between word reading and passage reading score. (E) Partial correlation of each factor with passage reading scores after regressing out all other factors. Asterisks denote significant correlation (* is p < 0.05, ** is p < 0.005, and so on). Error bars represent ± 1 s.d. of the correlation coefficient, calculated as in (D).

The part-sum model yielded excellent fits to the observed bigram dissimilarities (model correlation: r = 0.92 for upright bigrams, r = 0.94 for inverted bigrams; Figure 2B). Model correlations were close to the split-half consistency between participants, suggesting that the model explains nearly all the explainable variance in the bigram dissimilarities. Importantly, model fits were not systematically different between upright and inverted searches as would be expected if there were upright bigram detectors (Figure 2B). This in turn suggests that the better discrimination of upright bigrams by participants must be driven by letter-level differences in the part-sum model parameters.

We obtained several interesting insights upon a deeper investigation of the part-sum model parameters. First, the single letter relations estimated by the part-sum model for the corresponding, across and within terms were correlated with the observed single letter dissimilarities in this experiment (r = 0.76, p < 0.005; r = 0.84, p < 0.0005 & −0.61, p < 0.05 for C, X & W terms, for the part-sum model fit to the average dissimilarities for upright bigrams across all participants). Second, the within-bigram terms are consistently negative (Figure 2C), suggesting that search is harder when bigrams contain dissimilar letters. We have observed this effect consistently in previous studies – it resembles the well-known finding that search is harder when distractors are heterogeneous (Duncan and Humphreys, 1989; Vighneshvel and Arun, 2013; Pramod and Arun, 2016). Third, the interaction between the letters (both across and within) were weaker for upright compared to inverted bigrams. This weaker interaction leads to improved search for upright letters by increasing their discriminability.

### Relation between bigram searches and reading fluency

The above findings that fluent readers are faster at discriminating upright bigrams might also be predicted by other covarying factors such as their RAN score, motor speed, overall executive function etc. To investigate these possibilities, we sought to predict the individual variation in reading fluency using a variety of possible factors. To avoid overfitting, we generated a predicted fluency score by training each factor on the word reading scores, and then compared this prediction with the passage reading score.

To characterize the effect of overall task performance for each subject, we included the motor speed (measured during a baseline motor block; see Methods) and overall accuracy (across all searches). To characterize any effects due to discrimination of single letters, we calculated the average dissimilarity across all single letter searches. To characterize the influence of upright bigrams, we fit a part-sum model to the upright bigram dissimilarities for each subject, and calculated the average of the corresponding, across and within terms separately, and included the constant term. We did likewise for the inverted bigram searches. Finally, we used the RAN score of each subject as a possible factor. For each factor we asked how well the predicted reading score using that factor matched the observed passage reading score.

The results of these analyses are summarized in Figure 2D. To establish an upper bound on model performance, we compared the word reading fluency and passage reading fluency scores, which were highly correlated (r = 0.91, p < 0.0005; Figure 1C). Among all the individual factors, the RAN score had the highest correlation with passage reading fluency (r = 0.55, p < 0.00005; Figure 2D), followed by the upright bigram terms (r = 0.40, p < 0.0005; Figure 2D). This correlation was best for the part-sum model terms, compared to other measures derived from the bigram searches (correlation of passage reading scores with average upright bigram dissimilarity of each subject: r = 0.32, p < 0.05; with the average difference between upright and inverted bigram dissimilarity: r = 0.36, p < 0.005). Thus, the part-sum model parameters seem to capture the essential aspects of bigram processing.

The above analysis shows that a number of factors are correlated with passage reading fluency, but there could be correlations between these factors. To assess the unique contribution of each factor, we performed a partial correlation analysis. Specifically, we asked whether the correlation between a given factor with the passage reading fluency score would continue to be significant after regressing out all other factors. This revealed only two factors with a significant partial correlation: upright bigram terms and RAN score (Figure 2E). Hence, we conclude that RAN and upright bigram terms uniquely predict reading fluency compared to all other factors.

## EXPERIMENT 2: BIGRAM SEARCHES WITH VARYING SPACING

The above findings show that reading fluency is associated with upright but not inverted bigram processing, suggesting that familiarity with upright letter orientations leads to specific changes in visual processing. We therefore wondered whether this effect would also be specific to frequently encountered letter spacings. This is an important question by itself because changes in letter spacing affect reading speed (Zorzi et al., 2012; van den Boer and Hakvoort, 2015; Hakvoort et al., 2017). In addition, by testing the same participants after ~10 months, we also asked whether improvements in reading fluency can be predicted from changes in bigram processing.

To this end, we recruited 65 children for Experiment 2, of whom 59 children had participated in Experiment 1 ~10 months earlier. Participants were again given the two reading tasks (word & passage reading), a RAN task, and a visual search task involving upright and inverted bigrams with normal or large spacing. All bigram searches were interleaved. An example bigram search array using normal letter spacing is shown in Figure 3A, and the same search with large spacing is shown in Figure 3B. It can be seen that the search with the large letter spacing is harder but this effect is weaker if the arrays are inverted. This was indeed true in general as well (see below).

**Figure 3.**
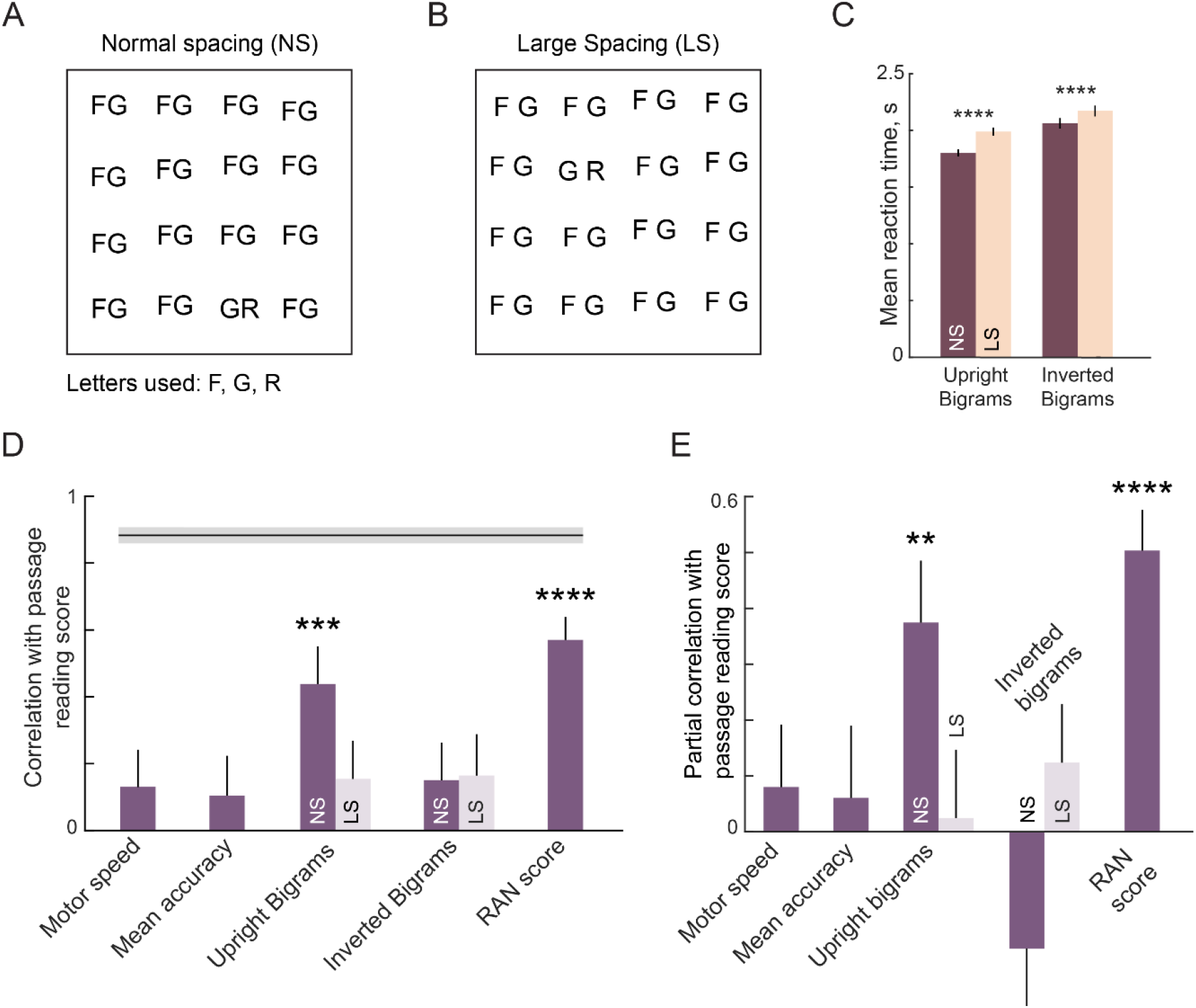
Effect of letter spacing on visual representation (Experiment 2) (A) Example upright bigram search array with small letter spacing. (B) Same as (A) but with large letter spacing. It can be seen that this search is slightly harder than the search in (A). (C) Average search times in the oddball search task for upright and inverted bigrams with normal and large spacing. Error bars indicate s.e.m. across participants. Asterisks denote statistical significance of the difference in means (**** is p < 0.00005, ANOVA – see text). (D) Model correlation of each factor predicting passage reading score. Error bars indicate ±1 s.d. using a bootstrap procedure, whereby we repeatedly sampled 67 participants with replacement for a total of 1,000 times. Shaded error bars on the top represents noise ceiling i.e. correlation between word reading and passage reading score. (E) Partial correlation of each factor with passage reading scores after regressing out all other factors. Asterisks denote significant correlation (* is p < 0.05, ** is p < 0.005, and so on). Error bars represent ± 1 s.d. of the correlation coefficient, calculated as in (A).

Overall, participants were highly accurate across all search types (accuracy, mean ± sem: 96% ± 0.5% for upright-normal spacing, 95% ± 0.5% for upright-large spacing; 95% ± 0.6% for inverted-normal spacing, 94% ± 0.7% for inverted-large spacing). They were also highly consistent in their responses (split-half correlation between RT of odd- and even-numbered participants, for normal and large letter spacing: r = 0.96 & 0.95 for upright bigrams, r = 0.96 & 0.97 for inverted bigrams; all p < 0.00005).

Participants responded significantly slower for upright bigrams with large spacing (average response times, mean ± sem across participants: 1.8 ± 0.03 s for normal spacing, 1.99 ± 0.04 s for large spacing; F(1, 8095) = 101.0, p < 0.00005 for main effect of spacing, ANOVA on RT with subject, spacing & image pair as factors; F(35, 8095) = 56.69, p < 0.00005 for image-pair, F(35, 8095) = 4.05, p < 0.00005 for interaction effect; Figure 3C). This effect was present even for inverted bigrams (average response times, mean ± sem across participants: 2.06 ± 0.05 s for normal spacing, 2.17 ± 0.05 s for large spacing, F(1, 8095) = 26.4, p < 0.00005 for main effect of spacing, ANOVA on RT with subject, spacing & image pair as factors; F(35, 8095) = 64.12, p < 0.00005 for image-pair, F(35, 8095) = 2.05, p < 0.00005 for interaction effect).

The normal spacing advantage was larger for upright compared to inverted bigrams (average difference in RT between normal and large spacing searches, mean ± sem across participants: 0.19 ± 0.02 s for upright bigrams, 0.11 ± 0.02 s for inverted bigrams, p < 0.05 on a paired t-test across subject-wise differences). However, search dissimilarities were highly correlated with each other for both normal and large spacing searches (r = 0.94 for upright bigrams, r = 0.95 for inverted bigrams; p < 0.00005), as well as between upright and inverted conditions (r = 0.95 for normal spacing, r = 0.96 for large spacing; p < 0.00005). Thus, bigram dissimilarities are qualitatively similar across letter spacing and bigram orientation.

### Can reading fluency be predicted by bigram processing at the familiar spacing?

Next, we fit the part-sum model to the observed search dissimilarities for each subject for each of the four search types (upright/inverted x normal/large spacing). We then performed a similar analysis as before to determine whether the passage reading score can be predicted by various factors. The correlation of each factor with passage reading score is shown in Figure 3D. Interestingly, only the part-sum model terms for upright bigrams with normal spacing predicted reading fluency, compared to model terms for large spacing and inverted bigram terms (Figure 3D). As before, this correlation was specific to the part-sum model terms, compared to other measures from the bigram searches: passage reading fluency was only weakly correlated with the average upright bigram dissimilarity of each subject (r = 0.17 & 0.11 for small and large spacing, p = 0.19 & 0.37 respectively) and with the average difference between upright and inverted bigram dissimilarity (r = 0.02 & 0.03 for small and large spacing, p = 0.89 & 0.84 respectively). Thus the part-sum model captured some essential underlying aspect of bigram processing relevant to reading fluency.

To assess the unique contribution of each factor towards explaining reading fluency, we performed a partial correlation analysis as before. Only two factors showed a significant partial correlation with the passage reading score after regressing out all other factors: upright bigram terms for normal spacing and the RAN score (Figure 3E). Hence, we conclude that the effect of visual processing on reading fluency is highly specific both to the familiar (upright) orientation and familiar (normal) spacing.

### Can bigram processing changes predict longitudinal changes in fluency?

Since the same participants were tested ~10 months apart in Experiments 1 & 2, we wondered whether improvements in reading fluency can be predicted using changes in upright bigram processing. We first compared the reading and RAN scores across Experiments. As expected, all scores improved with time (Figure 4A). To assess whether the change in reading scores can be predicted using the change in bigram processing, we took the difference in the average model term magnitudes of each type (corresponding, across, within, and constant terms) and asked whether the change in fluency can be predicted using a linear sum of the change in the model parameters for upright or inverted bigrams. We found that upright bigram terms were able to predict the improvement in both word reading and passage reading (r = 0.42, p < 0.005 for word reading, r = 0.29, p < 0.05 for passage reading; Figure 4B). By contrast, changes in inverted bigram processing predicted word reading only weakly (r = 0.30, p < 0.05; Figure 4B) but did not predict passage reading (r = 0.15, p = 0.27). Thus, only upright bigram processing changes robustly predicted fluency improvements.

**Figure 4.**
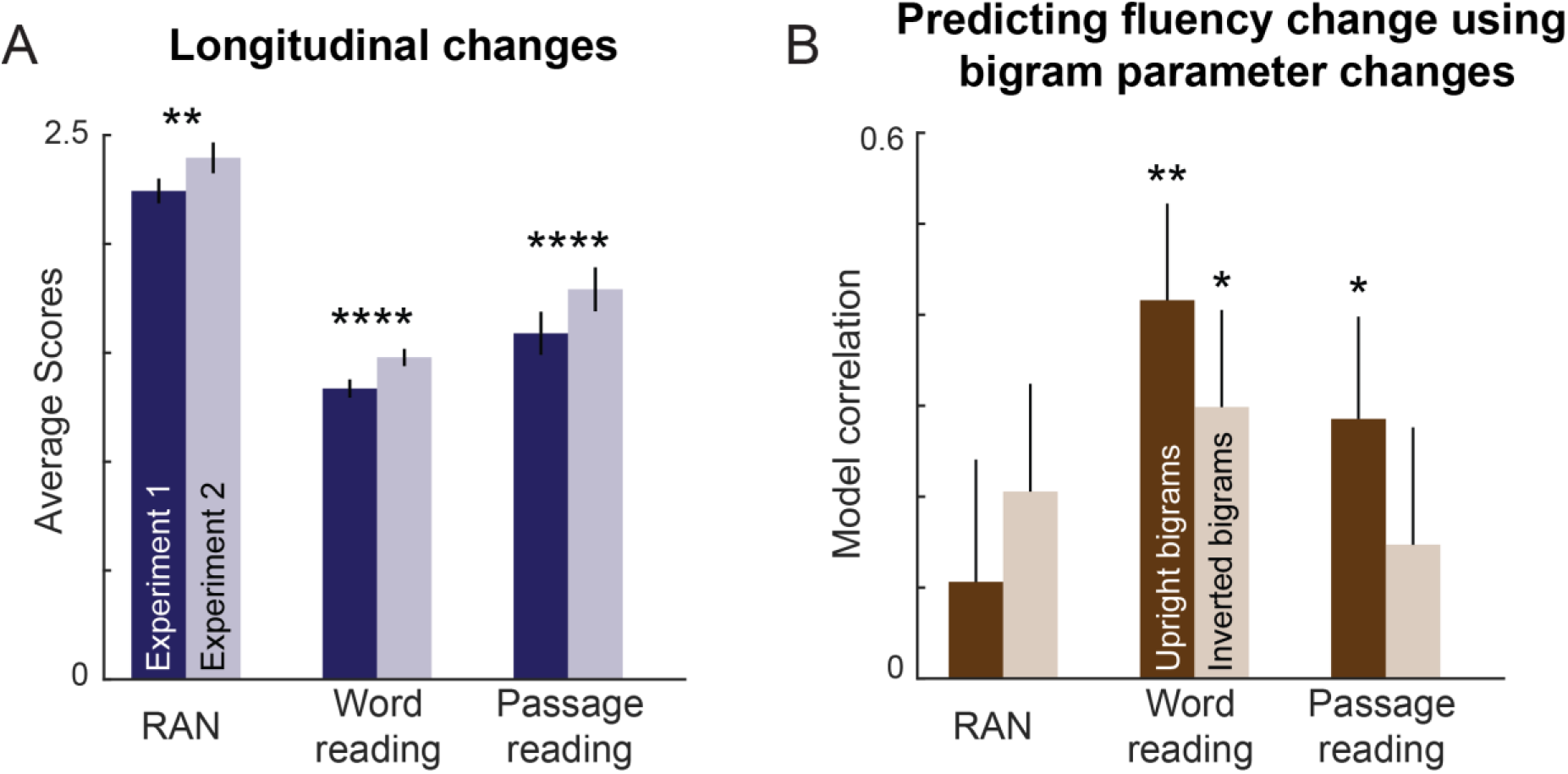
Longitudinal prediction of reading fluency using upright bigrams. (A) Change in fluency scores across different fluency measures with reading expertise. Asterisk represents statistical significance calculated using sign-rank test. Error bars represent s.e.m across participants. (B) Correlation between change in fluency scores with change in visual representation for upright (*dark*) and inverted (*light*) bigrams. Error bars represent indicate ±1 s.d. obtained by a bootstrap procedure, whereby we repeatedly sampled 59 participants with replacement for a total of 1,000 times. Asterisks denote statistical significance of each correlation (* is p < 0.05, ** is p < 0.005, and so on).

We conclude that longitudinal changes in reading fluency can be predicted using changes in upright bigram processing.

## DISCUSSION

Here we investigated whether reading fluency in children is associated with their performance on visual search tasks. Our main finding is that visual search for bigrams predicts reading fluency; this is only for upright (but not inverted) bigrams and with normal (but not large) spacing. This association predicted both cross-sectional inter-individual variations in reading fluency as well as longitudinal changes within individuals. Below we discuss these findings in relation to the existing literature.

We have found that reading fluency has a highly specific association with upright, normally spaced bigrams during visual search. This finding is consistent with crowding as well as serial position effects being different for letters compared to unfamiliar symbols (Grainger et al., 2010; Chanceaux and Grainger, 2012). It is also consistent with the processing deficits for letters but not symbols in dyslexic readers (Shovman and Ahissar, 2006). But the specificity of the association to bigrams in upright orientation with normal spacing is noteworthy, because such selective effects have not been reported previously. It suggests that visual representations for letters and bigrams undergo changes and these changes are specific to the orientation and spacing of text that is commonly encountered. It also indicates a possible resolution to conflicting evidence in the literature with regard to letter spacing. Some studies have found improved reading speed and accuracy with increased letter spacing (Zorzi et al., 2012; Hakvoort et al., 2017), whereas others have found that reading speed is optimal at the default spacing (Perea et al., 2011; van den Boer and Hakvoort, 2015). We speculate that these discrepancies could reflect differences in the statistics of letter characteristics (e.g., font, spacing, size) as experienced by sampled readers in different studies.

Our findings show an association between upright bigram processing and fluent reading, but do not reveal the direction of causality: does fluent reading lead to upright bigram processing, or does bigram processing lead to fluent reading? This question can be resolved if early changes in bigram processing were observed to precede changes in fluent reading, but this will require an extensive longitudinal study starting when literacy is emergent and while controlling for a number of other confounding factors. Nonetheless our findings do suggest a possible component in an intervention, whereby visual search activities involving upright bigrams or longer strings could facilitate optimal letter processing prior to the conversion of letters and letter strings into sounds and eventually words and their meaning.

Our results also reveal how visual representations change with reading. We have found that bigram discrimination in visual search can be explained entirely using dissimilarities between pairs of letters, for both upright and inverted bigrams. These results challenge the widely held view that reading should lead to the formation of specialized bigram detectors (Grainger and Whitney, 2004; Dehaene et al., 2005). If bigram detectors were formed through exposure to upright letters, upright bigram discrimination should have been less predictable from single letters compared to inverted bigram discrimination, but we observed no such trend (Figure 2B). Rather, we found that upright bigrams are more discriminable because of weaker within-bigram interactions (Figure 2C). We propose that reading not only makes single letters more discriminable but also makes letters more independent within a bigram, enabling the parallel processing of letters in a word.

We have found that RAN scores and upright bigram processing explained unique components of variance in reading fluency (Figure 2E, 3E). This is consistent with theoretical accounts of RAN that suggest it captures domain-general speed of processing (Kail et al., 1999; Sideridis et al., 2016), domain specific speed of access to phonological codes and visual features (Stainthorp et al., 2010), cross-modal print processing (Nag and Snowling, 2012) and recognition of whole items (Lervåg and Hulme, 2009). However our results go further to show that there are bigram-level changes in visual processing that also seem to enable reading fluency that are not captured by the single letter or digit naming processes integral to RAN. We speculate that the upright bigram processing measured in our study captures key aspects of orthographic processing that can complement other measures (RAN, phoneme awareness, executive function tests) to track the development of typical or atypical reading skills (Norton and Wolf, 2012).

## METHODS

All children and their parents/guardians gave informed consent to an experimental protocol approved by the Institutional Human Ethics Committee of Indian Institute of Science, University of Oxford and The Promise Foundation. All participants were students of a school in Bengaluru where English is the medium of instruction. All participants had normal or corrected to normal vision.

In both Experiments 1 & 2, participants were asked to perform two reading tasks (word reading and passage reading), a RAN task and a visual search task. The sample sizes were chosen based on previous studies in the literature and this age range was chosen because at this age there is broad individual variation in reading fluency. The reading and naming tasks were identical in both experiments and are summarized below.

### Reading & RAN tasks (Experiments 1 & 2)

#### Word reading task

This was the standardized sight word efficiency task (TOWRE). In this 104-word list, words increased in difficulty level, from simple words like “up” and “cat” to difficult words like “information” and “boisterous”. The word reading score was calculated as the number of words read correctly in the first 45 seconds, converted into a words/minute score.

#### Passage reading task

Participants were asked to read aloud a five line passage titled “Qasim’s kurta” describing the patterned dress of a stranger (Nag and Arulmani, 2015). The passage was edited to a word count of fifty. Participants were informed that they will have to answer two questions at the end of the passage and therefore had to read carefully. A discontinuation rule was applied after errors on eight words (an error rate of 15%). The passage reading score was calculated as the total number of words read correctly divided by the time taken up to the point attempted, in units of words/minute.

#### Rapid Automatized Naming (RAN)

A set of 40 digits arranged in a 5 x 8 grid was shown to the subject, which they had to read aloud. The RAN score was calculated as 40 divided by the time taken by participants to complete reading the digits.

### Experiment 1: Single letter and bigrams searches

#### Procedure

Participants were seated comfortably in front of a laptop monitor placed ~60 cm away under the control of custom programs written in HTML/Javascript.

#### Participants

A total of 68 children (34 male, aged 9.5 ± 0.9 years; 23 from 3^rd^ grade, 27 from 4^th^ grade, 18 from 5^th^ grade) were recruited for the study. One subject was excluded from the analyses due to the overall accuracy being less than 80%.

#### Stimuli

A total of 13 uppercase English letters (A, H, I, J, K, L, N, R, S, T, U, V, Y) were chosen for the single letter search task. These letters were chosen to contain similar and dissimilar letters. All letters were shown in the Arial Font with the exception of the letter ‘I’, for which horizontal bars were added at the top and bottom to improve its discriminability. The height of each letter was 1° in visual angle.

For the bigram task, 6 letters (A, L, R, S, T, and V) were combined in all possible manner (i.e. AA, AL, AR, AS, AT, AV, LA, LL, … etc) to form 36 bigrams. These letters were chosen because they were not symmetric along the horizontal axis. Inverted bigrams were created by flipping the upright bigrams.

#### Behavioural tasks

To ensure familiarity with the buttons and measure their motor speed, participants first performed a baseline block prior to visual search. In this block, a white circle appeared on either side of a vertical red line dividing the screen (10 trials) and participants responded its location using the same keys. The baseline block was followed by a practice block of visual search using unrelated objects (20 trials) and then followed by the main visual search block.

In the main visual search block, participants performed a total of 616 correct trials (^13^C_2_ = 78 single letter searches +115 upright bigram searches + 115 inverted bigram search and 2 repeats of each). We selected 115 searches out of 630 (^36^C_2_) possible searches to ensure a range of search difficulty. There were a total 15 pairs where first letter changes, 13 pairs where second letter changes, and 87 pairs with both letter changes. These 115 search pairs were fixed across all participants. All trials were interleaved, and incorrect/missed trials appeared randomly later in the task but were not analyzed.

The MATLAB function “isoutlier” was used to remove any data points that lie three scaled deviations away from the median. This was done to improve the split-half consistency of the data. We obtained qualitatively similar results without this step.

##### Part-sum model to explain bigram dissimilarities using single letters

For each of the 115 bigram searches, we calculated the average search time (averaged across repeats and participants) and converted this into search dissimilarity by taking the reciprocal (1/RT). This was done because previous work has shown that the reciprocal of search time yields better models of visual search compared to models based directly on RT (Arun, 2012; Pramod and Arun, 2014, 2016). According to the part-sum model, the net dissimilarity between two bigrams AB & CD is given by a sum of pairwise letter relations between letters at corresponding and opposite locations across bigrams and within-bigram relations. Specifically,

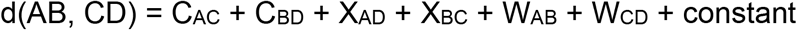

where C_AC_ & C_BD_ represent dissimilarity between letters at the corresponding locations of the two bigrams, X_AD_ & X_BC_ represent the dissimilarity between letters at opposite locations in the two bigrams, and W_AB_ & W_CD_ represent dissimilarity between letters within the two bigrams. This is a very general model because it allows for potentially different single letter dissimilarities of each type. It works because a given letter pair at each location can occur repeatedly across multiple bigram pairs (e.g. letter pair A-C is present at the corresponding locations of the pairs **A**B-**C**D, **A**D-**C**D, B**A**-B**C** etc.). Since bigrams were made from 6 possible letters, there are ^6^C_2_ (= 15) letter pairs for each of the corresponding, across, and within terms. This results in a 46-parameter model (15 letter pairs/term x 3 terms + 1 constant). Since we have 115 dissimilarities values and only 46 parameters, we can uniquely estimate all the parameters using linear regression. The resulting set of simultaneous equations can be represented as **y** = **Xb**, where **y** is a 115×1 vector of observed dissimilarities, **X** is a 115 x 46 matrix with entries of either 0, 1 or 2 depending on whether a particular pair is absent, present or repeated at each of the corresponding, across or within terms and **b** is a 46 x 1 vector of unknown weights.

To compare model parameters for upright and inverted bigrams (Figure 2), we fit a single model for both upright and inverted bigrams together with separate C, X, W terms for each orientation but a single constant term. To predict fluency scores for each subject (Figures 2 & 3), we fit the part sum model to upright and inverted dissimilarities separately.

##### Modelling fluency scores

For each subject, we estimated various factors from visual search experiment that could potentially predict reading fluency such as baseline reaction time, mean accuracy, mean single letter dissimilarities, part-sum model parameters estimated by modelling dissimilarities observed from upright and inverted bigram searches, and RAN score. To estimate the cross-validated fluency model fits, we trained each factor on word reading score and evaluated it against the passage reading score.

For each scalar factor, we fitted a linear model **y** = **Xb**, Here, **y** is a 67×1 vector of observed word reading score, **X** is a 67×2 matrix with entries containing one of the above mentioned factor along with a constant term, **b** is a 2×1 vector of unknown weights that are estimated after solving the linear regression (*regress* function in MATLAB). Next, we calculated the predicted reading score using the estimated weights i.e. 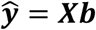 and correlated it with the passage reading score. The correlation coefficient quantifies the contribution of each factor in predicting reading fluency.

Since upright and inverted bigram factors contain multiple part-sum model parameters, we first averaged the estimated corresponding, across and within term interactions across all 15 letter pairs. This resulted in 4 parameters for each subject (including the constant term of the part-sum model). Next, we performed the same model fits as mentioned above to predict the fluency score as a linear combination of average model terms i.e. **y** = **Xb**,. Here, **y** is a 67×1 vector of observed word reading score, **X** is a 67×5 matrix with entries containing the average model terms together with a constant term, and **b** is a 5×1 vector of unknown weights.

##### Partial correlation analyses

To estimate the unique contribution of each factor, we performed a partial correlation analysis. First, we took the predicted fluency score for each factor (as described above) and regressed out the net contribution of all the other factors. Specifically, we fit a linear model **y = Xb**, where **y** is a 67×1 vector of fluency score predictions using that factor, and **X** is a 67-row matrix containing all the other factors, and **b** is a vector of unknown weights. We then calculated the residuals of this model i.e. (y – Xb) which represent the predictions of that factor that are not explained by the other factors. Proceeding likewise, we regressed out the net contribution of all the factors from the passage fluency score. The partial correlation is the correlation between these two sets of residuals, and represents the correlation between reading fluency and a particular factor that remains even after removing the influence of all other confounding factors.

### Experiment 2: Effect of letter spacing

All details of Experiment 2 were identical to those in Experiment 1 except those outlined below.

#### Participants

A total of 65 children (31 male, aged 10.2 ± 0.9 years, 23 from 4^th^ grade, 26 from 5^th^ grade and 16 from 6^th^ grade) were recruited 10 months later for this follow-up experiment. Of these 59 children had previously participated in Experiment 1.

#### Stimuli

A total of 3 letters (F, G, and R) were combined in all possible ways (i.e. FF, FG, FR, GF, … etc) to form a total of 9 bigrams. These letters were chosen because they were not symmetric along the horizontal axis. Letters were 1° in height, and were separated by either 0.18° (normal spacing) or 1.05° (large spacing). The normal spacing here approximates the spacing between letters in Arial font but with a fixed width between letters.

#### Task

Participants performed a total of 288 searches (^9^C_2_ = 36 bigrams x normal and large letter spacing x 2 configurations x 2 repeats).

#### Part-sum model

Since there are only ^3^C_2_ = 3 letter relations each for the corresponding, across and within term, the part-sum model had only 10 free parameters, which were estimated from a total of 36 bigram dissimilarities.

### Longitudinal analysis

To this end, we analysed the data from 59 participants common to both Experiments 1 & 2. To predict the change in fluency score using the change in the average part-sum model parameters (averaged across ^3^C_2_ = 3 terms for corresponding, across, within terms, together with the constant term), we performed a linear regression to predict the change in fluency as a weighted sum of the part-sum model parameters. Specifically, we fitted a linear model **y = Xb**, where **y** is a 59×1 vector depicting difference in fluency score (i.e. Experiment 2 – Experiment 1 scores), **X** is a 59 x 5 matrix with rows containing the difference between each type of model term together with a global constant term, and **b** is a 5×1 vector of unknown weights that is estimated using standard linear regression (*regress* function in MATLAB).

## ACKNOWLEDGEMENTS

We are grateful to the children, their guardians and the staff at the Sandeepani Academy for Excellence for their participation, and to Laxmi Sutar, Sandra Beula, Pooja Shah, B. Kala and Sanjana Nagendra from The Promise Foundation for assistance with data collection. This work was supported by a Senior Fellowship (IA/S/17/1/503081) from the Wellcome Trust-DBT India Alliance to SPA, and the 2018 Trinity Term Department of Education Small Research Grant to SN.

## AUTHOR CONTRIBUTIONS

All authors contributed to the overall study design. AA, SPA & SN designed experiments, AA implemented the experiment and collected data, AA & SPA analyzed and interpreted data with inputs from KVSH & SN, and AA and SPA wrote the manuscript with inputs from KVSH & SN.

